# Machine Learning Analysis of Motor Evoked Potential Time Series to Predict Disability Progression in Multiple Sclerosis

**DOI:** 10.1101/772996

**Authors:** Jan Yperman, Thijs Becker, Dirk Valkenborg, Veronica Popescu, Niels Hellings, Bart Van Wijmeersch, Liesbet M Peeters

## Abstract

**Background:** Evoked potentials (EPs) are a measure of the conductivity of the central nervous system. They are used to monitor disease progression of multiple sclerosis patients. Previous studies only extracted a few variables from the EPs, which are often further condensed into a single variable: the EP score. We perform a machine learning analysis of motor EP that uses the whole time series, instead of a few variables, to predict disability progression after two years. Obtaining realistic performance estimates of this task has been difficult because of small data set sizes. We recently extracted a dataset of EPs from the Rehabiliation & MS Center in Overpelt, Belgium. Our data set is large enough to obtain, for the first time, a performance estimate on an independent test set containing different patients.

**Methods:** We extracted a large number of time series features from the motor EPs with the highly comparative time series analysis software package. Mutual information with the target and the Boruta method are used to find features which contain information not included in the features studied in the literature. We use random forests (RF) and logistic regression (LR) classifiers to predict disability progression after two years. Statistical significance of the performance increase when adding extra features is checked with the DeLong hypothesis test.

**Results:** Including extra time series features in motor EPs leads to a statistically significant improvement compared to using only the known features, although the effect is limited in magnitude (∆AUC = 0.02 for RF and ∆AUC = 0.05 for LR). RF with extra time series features obtains the best performance (AUC = 0.75 ± 0.07), which is good considering the limited number of biomarkers in the model. RF (a nonlinear classifier) outperforms LR (a linear classifier).

**Conclusions:** Using machine learning methods on EPs shows promising predictive performance. Using additional EP time series features beyond those already in use leads to a modest increase in performance. Larger datasets, preferably multi-center, are needed for further research. Given a large enough dataset, these models may be used to support clinicians in their decision making process regarding future treatment.

## 1 Background

Multiple sclerosis (MS) is an inflammatory demyelinating and degenerative chronic disease of the central nervous system, with symptoms depending on the disease type and the site of lesions. Ideally, MS should be featured by an individualized and intense clinical follow-up and treatment strategy. Typical MS symptoms include sensation deficits and motor, autonomic and neurocognitive dysfunction, but the clinical course of MS varies greatly between individuals [1]. Establishing the prognosis for an individual is a major epidemiological goal in MS research. For the time being however, it remains impossible to accurately predict the disease course of an individual patient. This causes anxiety and frustration for patients, families and health-care professionals [2].

There are numerous measures available to study MS, but there is no gold standard in monitoring disease activity, measuring progression or evaluating short- and long term therapy efficacy. Besides magnetic resonance imaging (MRI) scans, which visualize lesions in the central nervous system, other clinical parameters are used in the assessment of MS disease progression [3, 4, 5, 6, 7]. The clinically most commonly used is the expanded disability status scale (EDSS) [8], ranging from 0 to 10 and quantifying the level of disability in MS patients. However, the potential limitations of EDSS have received increasing attention. Indeed, EDSS depends on the interpretation of the neurologist, is insufficiently sensitive to detect robust changes in disability of short time frames, and shows poor responsiveness to disease progression and treatment effects in secondary progressive (SPMS) and primary progressive MS (PPMS) [8].

Several research groups have shown that evoked potentials (EP) allow monitoring of MS disability, both in cross-sectional studies and longitudinal studies [9, 10, 11, 12, 13, 14, 15, 16, 17, 18, 19, 20, 21, 22, 23, 24, 25, 26, 27, 28, 29, 30, 31], see [32, 33] for reviews. EP provide quantitative information on the functional integrity of well-defined pathways of the central nervous system, and reveal early infra-clinical lesions. They have a predictive value regarding the evolution of disability [10, 32, 12, 13, 16, 17, 27, 14, 18, 20, 26, 23]. EP measure the electrical activity of the brain in response to stimulation of specific nerve pathways or, conversely, the electrical activity in specific nerve pathways in response to stimulation of the brain. Different types of EP are available corresponding to different parts of the nervous system [34]. For visual EP (VEP) the visual system is excited and conductivity is measured in the optic nerve; for motor EP (MEP) the motor cortex is excited and conductivity is measured in the feet or hands; for somatosensory EP (SEP) the somatosensory system (touch) is excited and conductivity is measured in the brain; and for brainstem auditory EP (BAEP) the auditory system (ears) is excited and conductivity is measured at the auditory cortex. EP are able to detect the reduction in electrical conduction caused by damage (demyelination) along these pathways even when the change is too subtle to be noticed by the person or to translate into clinical symptoms.

If several types of EP are available for the same patient this is referred to as a multimodal EP (mmEP). Considerable community effort has been performed to summarize mmEP by a one-dimensional statistic, called the EP score (EPS), by applying different scoring methods [13, 14, 17, 18, 30, 22, 31]. The scoring methods described in literature use a limited amount of features from these EP time series (EPTS). The latency (time for the signal to arrive) is always included. Besides latency, amplitude and dispersion pattern are also possibly included in the EPS [22]. Latencies and the EPS have been used as a biomarker in clinical and observational studies [35, 36, 37, 38, 39]. By only using two or three variables extracted from the EPTS, possibly useful information is lost. In this study, we investigate whether a machine learning approach that includes extra features from the EPTS can increase the predictive performance of EP in MS.

In literature, the main modeling techniques are linear correlation of latency or EPS with EDSS, and linear or logistic regression models. The performance of these linear models is most often measured by the explained variance (*R*^2^), mean-squared error (MSE), and area under the curve (AUC) of the receiver-operating characteristic (ROC). Except for one study with 30 patients [12], no study has used an independent test set to asses model performance. Some studies use cross-validation to estimate model performance [16, 18, 19, 26, 21]. Akaike information criterion (AIC) or Bayesian information criterion (BIC) are sometimes included to encourage model parsimony. While such models are statistically rigorous, insightful, and often used in practice [40], a realistic performance estimate is obtained by training on a large dataset (part of which is used as a validation set to tweak any hyperparameters), and testing on an independent large dataset containing different patients. This study provides, for the first time, such a performance estimate.

Because of its unpredictable disease course, most clinical interest lies in predicting disease progression. This is often translated to a binary problem, where a certain increase in EDSS is considered as a deteriorated patient. In the literature, AUC values for this task range from 0.74 to 0.89, with prediction windows between 6 months and 20 years [19, 26, 22, 23, 25].

We recently extracted a large number of EPTS from the Rehabiliation & MS Center in Overpelt, Belgium. This patient cohort consists of individuals undergoing treatment. Despite the fact that this adds another unknown to the problem, due to the incomplete nature of the treatment records, it is, clinically speaking, the most relevant scenario. In a clinical setting, the majority of patients will have had some form of treatment prior to these types of measurements. The resulting dataset, containing the full time series of mmEP with longitudinal information for most patients, is the first of its kind. A description of the dataset and how to access it will be provided in a separate publication [41]. We perform a disability prediction analysis on the MEP from this dataset, as this EP modality is most abundant in the dataset. A machine learning approach is used to see if there is extra information in the MEP for predicting disability progression after 2 years, besides latency and amplitude. This prediction of progression can be used to support a clinician’s decision making process regarding further treatment. 419 patients have at least one measurement point, where previous studies had between 22 and 221 patients. Including extra EP features leads to a statistically significant increase in performance in predicting disability progression, although the absolute effect is small. Our results suggest that this effect will become more stable on a larger dataset. We show that a nonlinear model (random forests) achieves significantly better performance compared to a linear one (logistic regression).

## 2 Methods

### 2.1 Dataset

The full evoked potential dataset consists of 642 patients and has SEP (528), BAEP (1526), VEP (2482), and MEP (6219) visits [41]. We only study the MEP, because they are most frequently measured. Each MEP visit contains 4 measurements: two for the hands (abductor pollicis brevis (APB) muscle), and two for the feet (abductor hallucis (AH) muscle). Visits that don’t contain all 4 EPTS are discarded. An example of the EPTS for a single visit is shown in Figure 1. We use the standard definition of disability progression [42], where the patient has progressed if EDSS_*T*_1__ − EDSS_*T*_0__ >= 1.0 for EDSS_*T*_0__ ≤ 5.5, or if EDSS_*T*_1__ − EDSS_*T*_0__ >= 0.5 for EDSS_*T*_0__ > 5.5. T_0_ is the time of the first measurement, and T_1_ is the time of the EDSS measurement between 1.5 and 3 years which is closest to the 2 year mark. The MEP visit has to occur 1 year before or after T_0_. Visits without two-year follow-up are discarded.

**Figure 1.**
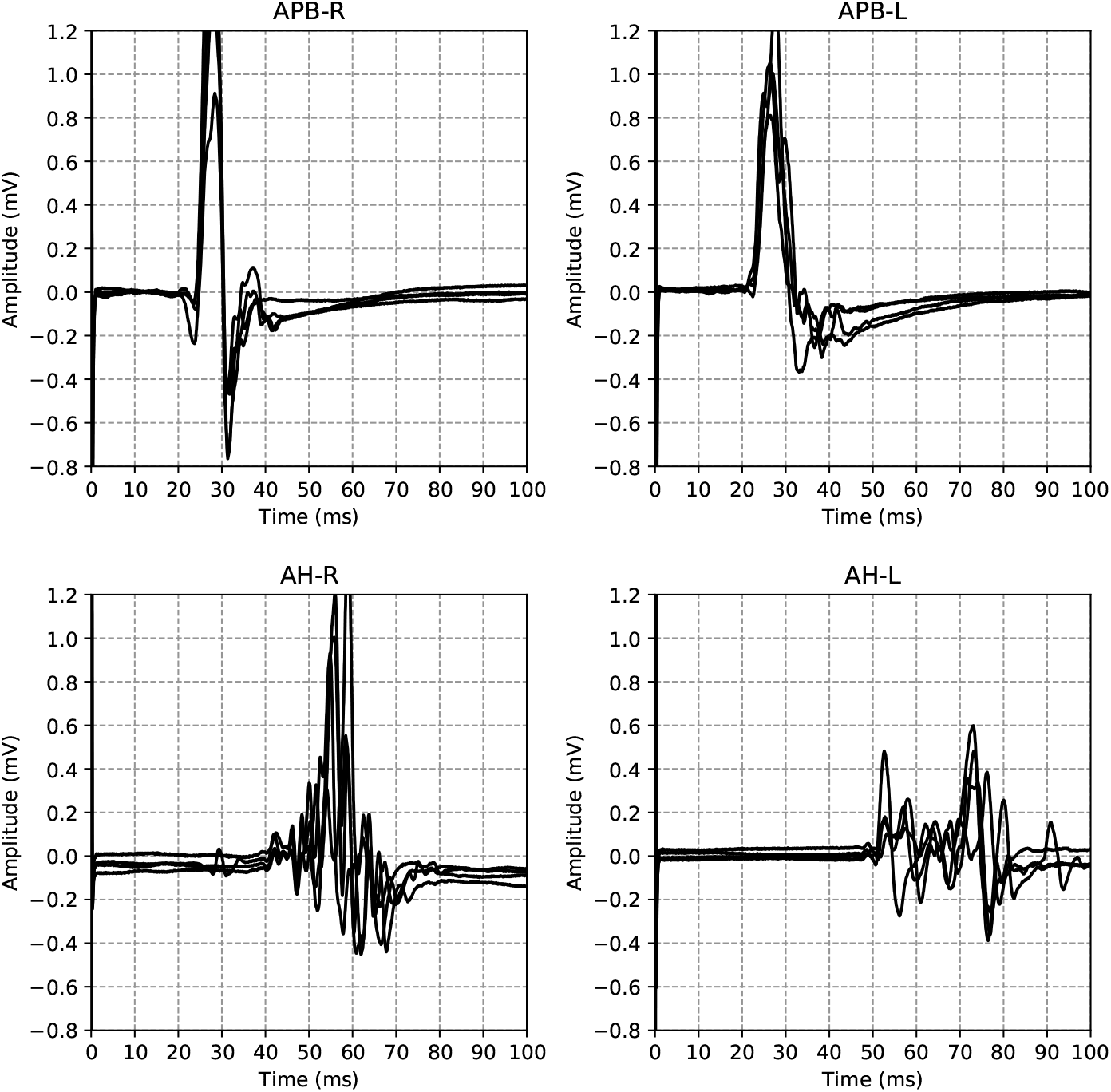
EPTS example. Example of the EPTS measured during a single patient visit. The titles indicate the anatomy (APB for the hands, AH for the feet) and the respective sides (L for left, R for right).

Measurements of a duration differing from 100ms are discarded. Around 97% of the data has duration 100ms, so this has little influence on the dataset size, and keeps the data homogeneous. The majority of EPTS consist of 1920 samples. Due to a slight difference in machine settings, some EPTS consist of 2000 samples. These EPTS were downsampled to 1920 samples.

In case no spontaneous response or MEP in rest position is obtainable, a light voluntary contraction of the muscle in question is asked in order to activate the motor cortex and increase the possibility of becoming a motor answer. This so called facilitation method is usually very noisy due to baseline contraction of the muscle measured, so we decided to drop them from the dataset altogether. Facilitated measurements are characterized by a non-flat signal right from the start of the measurement. We drop any EPTS that have a spectral power above an empirically determined threshold at the starting segment of the measurement. This segment is determined by the values of the latency of a healthy patient, which we set to be 17 ms as this is the lower bound for the hands. We use the same threshold for the feet, which is not a problem since the lower bound there is higher.

For each limb, the EP measurement is repeated multiple times. After discussion with the neurologists we decided to use only the EPTS with the maximal peak-to-peak amplitude, as this is likely to be the most informative measurement. The type of MS was inferred from the diagnosis date and the date of onset, both of which have missing values, making the type of MS field somewhat unreliable. The latencies are annotated by the medical staff for a subset of the time series.

After all these steps, we are left with a dataset of 10 008 EPTS from 2502 visits of 419 patients. Note that one patient can have several visits that satisfy the conditions for two-year follow-up. We have one target (worsened after 2 years or not) for each visit, so the total number of samples is 2502. Some of the characteristics of the dataset are summarized in Table 1.

**Table 1.** Characteristics of the dataset. The final column (% worsening) represents the percentage of visits of patients that have worsened 2 years later. Abbreviations used: MS multiple sclerosis, SD standard deviation, EDSS expanded disability status scale, F female, M male, CIS clinically isolated syndrome, PPMS primary progressive MS, RRMS relapsing-remitting MS, SPMS secondary progressive MS.

| MS type | # patients | age (SD) | EDSS (SD) | F / M ratio | # visits | % worsening |
| --- | --- | --- | --- | --- | --- | --- |
| Unknown | 138 | 44 (12) | 3.1 (2.0) | 98 / 40 | 558 | 13.1 |
| CIS | 7 | 31 (11) | 1.3 (0.8) | 7 / 0 | 9 | 11.1 |
| PPMS | 12 | 57 (11) | 4.0 (1.4) | 7 / 5 | 74 | 12.2 |
| RRMS | 223 | 44 (11) | 2.6 (1.5) | 164 / 59 | 1592 | 9.6 |
| SPMS | 45 | 56 (8) | 5.0 (1.5) | 31 / 14 | 269 | 14.5 |
| All | 419 | 45 (12) | 3.0 (1.8) | 301 / 118 | 2502 | 11.0 |

### 2.2 Data analysis pipeline

We start with a simple model that uses a subset of the features proposed in the literature. As other EPS require neurologist interpretation, and are therefore difficult to automate, we use latencies as a baseline. The fact that this is a fair baseline is supported by [22], where it was shown that different EPS have similar predictive performance, with short-term change or baseline values in (z-scored) latencies being more predictive than changes in other EPS. It is furthermore supported by the results from [16], where the central motor conduction time of the MEP was more informative for disability progression than the MEP EPS.

Despite the increased size of the dataset, the disability progression classification task remains a challenging problem. Challenging aspects are the limited sensitivity to change of the EDSS measure, its dependence on neurologist interpretation, and the heterogeneity of disease development. Therefore, our data analysis pipeline is mainly focused on minimizing overfitting. As our dataset includes the full EPTS, we wish to find one or more time series features that provide supplemental information on disability progression, on top of the features already used in the literature. A schematic overview of the data analysis pipeline is shown in Figure 2. The various steps in the data analysis pipeline are detailed below. The data analyis pipeline was implemented in Python using the scikit-learn library [43], with the exception of the Boruta processing step, for which we used the Boruta package in R [44], and the feature extraction, for which we used the highly comparative time-series analysis (HCTSA) package [45, 46] which is implemented in Matlab.

**Figure 2.**
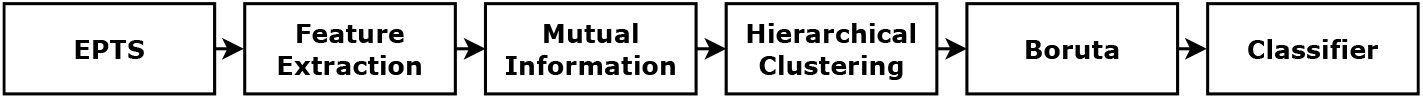
Pipeline. Schematic overview of the data analysis pipeline

#### Feature extraction

Because each EPTS starts with a large peak at the beginning, an uninformative artifact of the electrophysiological stimulation, the first 70 samples of each EPTS are discarded. A diverse and large set of time series features is extracted with the HCTSA package, which automatically calculates around 7700 features from different TS analysis methodologies. The motivation for this approach is to perform a wide variety of time series analysis types, and draw general conclusions on what approaches are useful. It makes the analysis less subjective, since one does not have to choose a priori the type of extracted features. Given the large size of this feature set, one expects that almost all useful statistical information contained in the EPTS is encoded in it. The feature matrix **F**_*ij*_ has rows *i* for each EPTS and columns *j* for each feature. If a column **f**_*j*_ contains an error or NaN value it is discarded. Normalization is performed by applying the following transformation on each column:

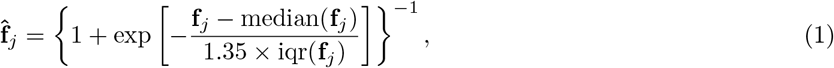

with iqr the interquartile range. Because the median and iqr are used, this normalization is robust to outliers. All normalized columns 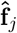 that contain an error or NaN are discarded. To exploit the symmetry between the measurements performed on the left and the right limb, we sum the features of both sides. This reduces the number of features we need to consider, which is helpful against overfitting. The final normalized feature matrices 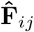 of AH and APB both have size 5004 *×* 5885.

#### Mutual Information

Our goal is to use a feature selection algorithm in order to determine the most important features. The ratio of the number of samples to the number of features is quite small (≈ 1). The feature selection algorithm we use, Boruta [44], was expected to work well for such a ratio [47]. We however found it to perform poorly for our problem. The performance of Boruta was tested by adding the latency, which is known to be relevant, to the list of candidates, which was subsequently not marked as relevant by Boruta. We therefore reduce the number of features using mutual information with the target as a measure of feature importance. We select the top ten percent of features based on this metric.

#### Hierarchical clustering

In this step we seek to reduce redundancy in our choice of features. We estimate this redundancy using the correlation distance, which we define here as

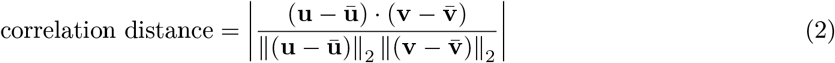

where **u** and **v** are the feature vectors we wish to compare, and ║⋅║_2_ the Euclidean distance. Note that we take the absolute value here so highly anti-correlated features are filtered as well. Features which are highly correlated have a distance close to zero, and conversely features which are not correlated have a distance close to 1. We cluster all features at a cutoff of 0.1 and keep only one feature for each cluster. In practice we found that this step eliminates few to no features.

#### Boruta

With the number of features now reduced to a more manageable count we run the Boruta algorithm [44] to estimate the importance of the remaining features. In a nutshell, the Boruta algorithm compares the importance (as determined by a Z-scored mean decrease accuracy measure in a random forest) of a given feature with a set of shuffled versions of that feature (called shadow features). If a feature’s importance is significantly higher than the maximal importance of its set of shadow features, it is marked as important. Conversely, any feature with importance significantly lower than the maximal importance of its shadow features is marked as unimportant, and is removed from further iterations. This procedure is repeated until all features are have an importance assigned to them, or until a maximal number of iterations is reached. We add a few literature features to the set of TS features as well (latencies, EDSS at *T*_0_ and age). There are multiple reasons for doing this. First off, it allows us to check the performance of the Boruta algorithm, as these features are known to be important. Secondly, some of the TS features may only be informative in conjunction with a given literature feature. Boruta returns a numerical measure of feature importance, which allows us to assign an ordering to the features. On average, some 80 features are confirmed to be relevant. From these we select the 10 most important ones, based on their importance score. This cutoff was chosen empirically using cross-validation, as more features leads to overfitting of the classifier.

#### Classifier

For the final classification we use a random forest, with 100 decision trees and balanced class weights. Using more trees led to no improvement in cross-validation. We opted for a random forest classifier due to the fact that this classifier is known to be robust against overfitting. We regularize the model further by setting the minimal number of samples required for a split to be 10% of the total number of samples. This value was obtained using cross-validation on the training set. The maximum depth of the resulting decision trees averages around 8. As linear models are often used in other works, we use logistic regression for comparison.

As discussed earlier, we have 4 time series per visit, 2 of the hands (left and right), and 2 of the feet (left and right). We run the pipeline for the hands and feet separately and average the predictions of the resulting classifiers to get the final prediction. This approach was chosen for two reasons: The time series resulting from the measurements are quite disparate, therefore the same time series features may not work well for both. The other reason is that adding too many features to the model causes the classifier to overfit. Splitting up the task like this reduces the number of features per model.

We found that the performance of the algorithm is greatly influenced by the choice of training and test set. To get a measure for how much this factors in we run this data analysis pipeline 1000 times, each time with a different choice of train/test split. That way we can get a better understanding of the usefulness of this process, rather than focusing on a single split. We ensure that patients don’t occur both in the training and the test set, and that the balance of the targets is roughly the same for the train and the test set. To illustrate, for some splits we actually obtain AUC values of 0.97, whereas others are random at 0.5. Of course, these are just the extreme values, the performance turns out to be normally distributed around the reported results. At this point, we have the ranking of the 10 most important features as determined by the Boruta algorithm. For the final prediction, we will add the top-*n* features. The value of *n* is determined on a validation set. For each train-test split, we use half of the test set as a validation set. This split into validation and test set satisfies the same conditions as before, and we evaluate the model for 100 such splits. So in total 1000 models are trained (on the training set), and are subsequently evaluated 100 times each, leading to 100 000 test set performances.

There is a trade-off to be taken into account. On the one hand we want as much data as possible to fit our model, which would require allocating as much data as possible to the training set. On the other hand, however, we want to accurately measure the performance of said model on an independent test set, which for a heterogeneous dataset also requires a large amount of data to minimize the variance. To get an idea of both extremes we evaluate the pipeline at various splits of the dataset. We run the entire pipeline for 4 different sizes of the training set, composed of 20, 30, 50 and 80 percent of the dataset.

## 3 Results

### 3.1 Disability progression task

Here we present the results of the disability progression prediction task. In the literature, the main features that are considered are: Latency, EDSS at *T*_0_, peak-to-peak amplitude, age, gender and type of MS. We note that not all of these are found to be significant in the literature (see, e.g., [25]). Using cross-validation we determined that using the latencies of the left and right side separately, the EDSS at *T*_0_ and the age worked best for this prediction task. Adding additional literature features leads to a negligible performance increase. We assess the performance of the literature features as well as the performance when we add additional time series features.

The main results are shown in Figure 3. As is to be expected, we see that the overall performance of the pipeline increases as the size of the training set increases, while the variance of the result also increases due to the smaller size of the test set. The general trend we see is that adding the extra time series features improves the performance on the independent test set, but only marginally. RF performs better than LR both with and without the additional TS features, with the difference being especially evident when not adding them. The figure indicates that increasing the dataset size further would improve the performance.

**Figure 3.**
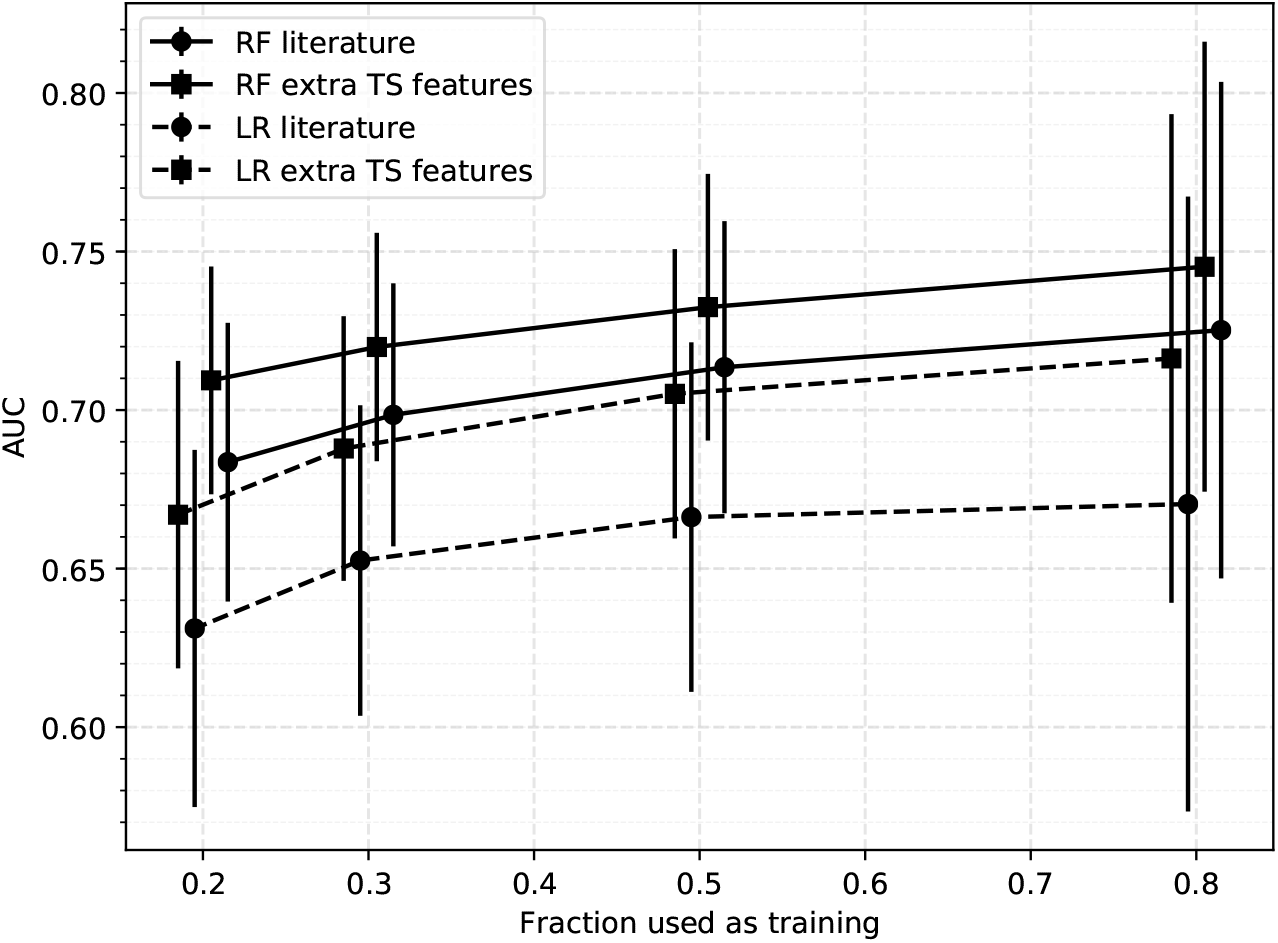
Results of the disability progression task. Results are shown for different sizes of training set. Each point represents an average over 100 000 test sets, with the error bar indicating the standard deviation. Results are shown for the baseline model which uses a subset of known features (Latency, EDSS at *T*_0_ and age), as well as a model where we add additional TS features. Abbreviations used: RF Random Forest, LR Logistic Regression, TS Time series. These results are represented numerically in Table 2.

As a check for our assumption of using only a subset of the literature features, we also checked the performance when adding additional literature features to the classifier (peak-to-peak amplitude, gender and type of MS). The resulting model performed worse than the model using just 4 literature features in almost every case, and in the cases where it does increase it does so by a negligible margin. It also degrades the performance gain by adding TS features, presumably due to overfitting. This reaffirms our decision of using just the latencies, the age and the EDSS at *T*_0_.

**Table 2.**
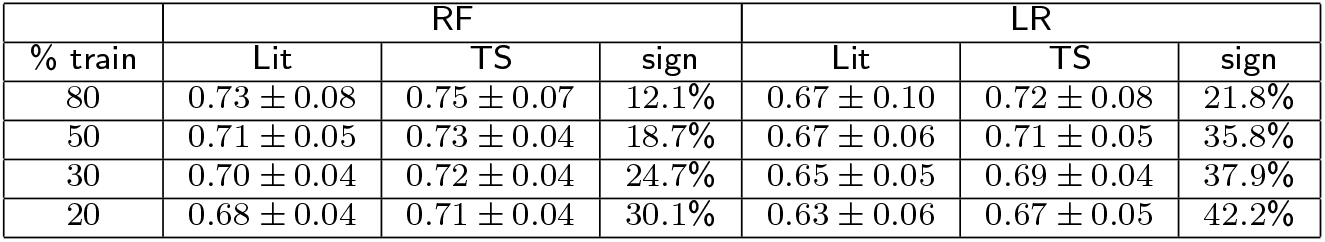
Results of the disability progression task The leftmost column indicates what percentage of the dataset was used for training. Results are shown for the classifier using just latencies, EDSS at *T*_0_ and age (Lit), and for the classifier trained on these features + additional TS features (TS). RF (Random Forest) and LR (Logistic Regression) indicate the classifier that was used. The *sign* column indicates the percentage of splits with a significant improvement, according to the DeLong test. These results are shown graphically in Figure 3.

| % train | RF |  |  | LR |  |  |
| --- | --- | --- | --- | --- | --- | --- |
|  | Lit | TS | sign | Lit | TS | sign |
| 80 | $0.73 \pm 0.08$ | $0.75 \pm 0.07$ | 12.1% | $0.67 \pm 0.10$ | $0.72 \pm 0.08$ | 21.8% |
| 50 | $0.71 \pm 0.05$ | $0.73 \pm 0.04$ | 18.7% | $0.67 \pm 0.06$ | $0.71 \pm 0.05$ | 35.8% |
| 30 | $0.70 \pm 0.04$ | $0.72 \pm 0.04$ | 24.7% | $0.65 \pm 0.05$ | $0.69 \pm 0.04$ | 37.9% |
| 20 | $0.68 \pm 0.04$ | $0.71 \pm 0.04$ | 30.1% | $0.63 \pm 0.06$ | $0.67 \pm 0.05$ | 42.2% |

### 3.2 Significance test of performance increase

To check whether the increase in performance by adding TS features is significant, we employ the DeLong test [48] which tests the hypothesis that the true difference in AUC of the model with and without TS features is greater than zero. For each split we compare the ROC curves of the classifier with and without the additional TS features. The results are shown in Figure 4. We observe that the percentage of splits with significantly improved performance increases with the size of the testset, reaching a maximum at 80% of the dataset used for the testset. We argue that the low fraction of significant improvement is mainly due to the power of the test. To support this further we show the significance percentages for a single model (the one trained on 20% of the dataset), tested on subsets of the remaining 80% of increasing size. The results are shown in Figure 5, from which we see the fraction of significant splits increases steadily with the number of samples in the test set.

**Figure 4.**
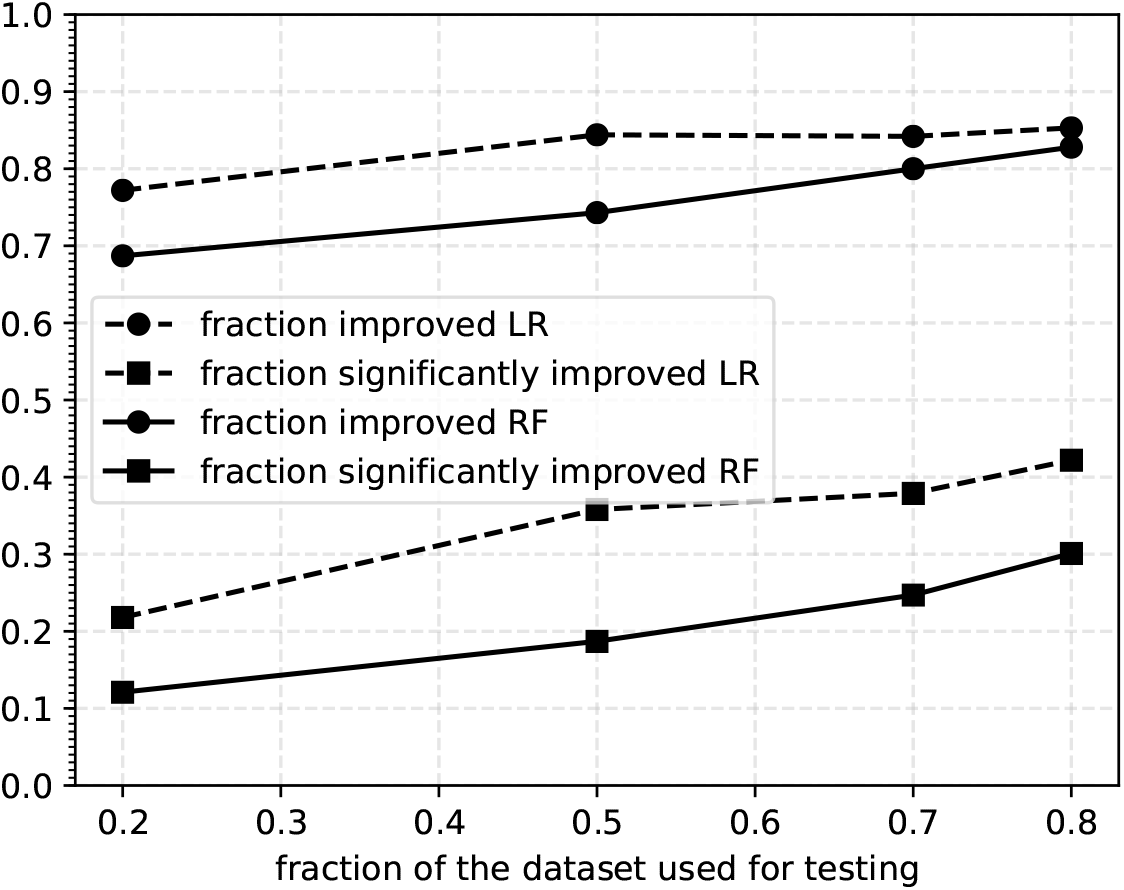
Significance results. The results of the DeLong test on the improvement by adding TS features. We show both the fraction of splits that show an improvement and the fraction of splits that show significant improvement according to the DeLong test.

**Figure 5.**
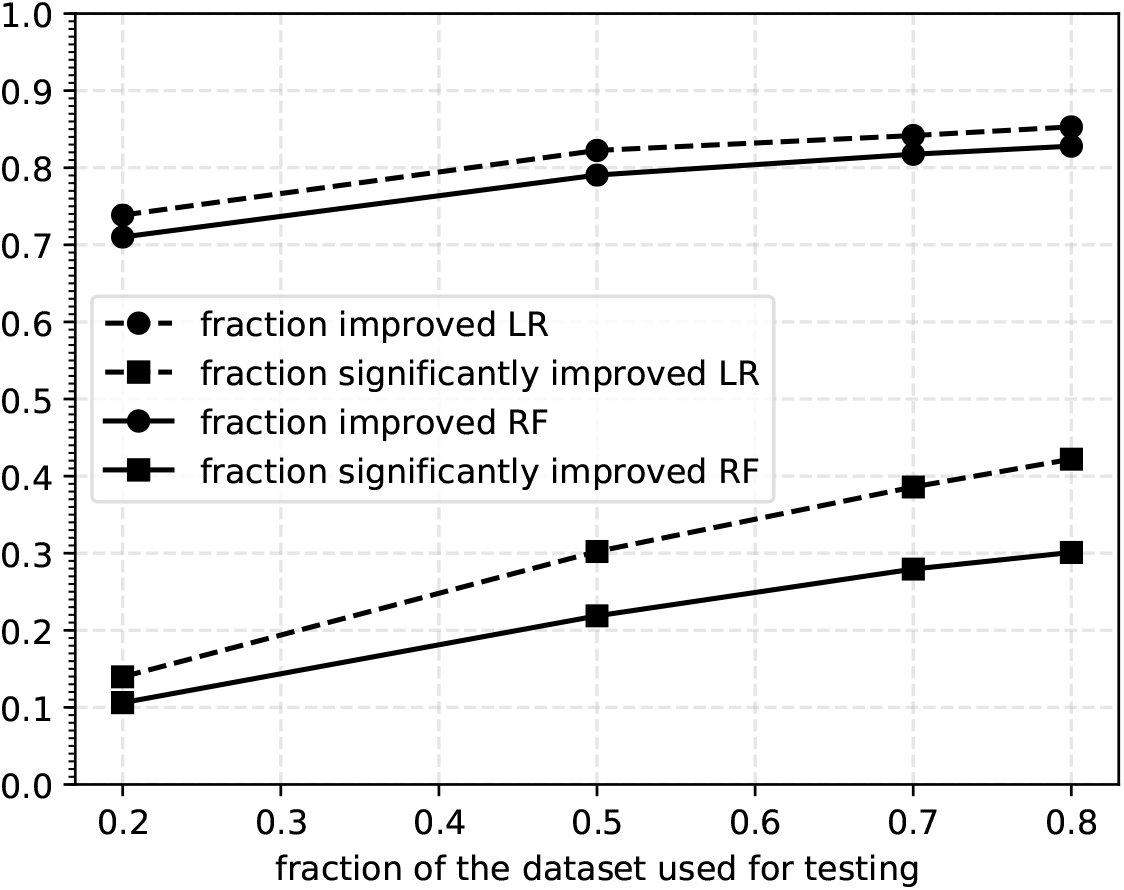
Significance results single model. Results of the significance tests, using a single model (trained on 20% of the dataset), and tested on various sizes of test set. Both the fraction of improved splits and the fraction of significantly improved splits are shown. The trend suggests more data would likely increase the fraction of splits that show significant improvement.

### 3.3 Selected features

It is interesting to see which TS features are often found to be important according to our feature selection method. As the pipeline is run independently for 1000 times we have 1000 ranked sets of features deemed important by the feature selection. We consider only the train/test-split where the training set consists of 80% of the dataset, as the feature selection is most stable in this case. We consider the anatomies separately as the selected TS features are different for each. Here we give only a brief overview of the features that we found to be most important. For a ranked list of the 20 most important features for both APB and AH we refer the reader to the additional files [see Additional file 1]. There we also provide a way of obtaining the code used to generate these features.

#### APB

The feature most often found to be important ranks in the final 10 features for 83.9% of splits. In 74.7% of splits it ranks in the top 3. The feature in question is calculated by sliding a window of half the length of the TS across the TS in steps of 25% of the TS (so a total of 3 windows is considered). For each window, the mean is calculated. Finally, the standard deviation of these means, divided by the standard deviation of the entire TS, is calculated. In practice this feature seems to characterize how fast the TS returns to an average of zero after the initial peak. The other high-ranking features are mostly other sliding window calculations or features that compute characteristics of the spectral power of the TS. The prominence of these features drops off quickly, e.g., the second highest ranking feature occurs in the top 4 for 39% of splits.

#### AH

For AH one feature in particular stands out. It is included in the final 10 features for 97.5% of splits. In 90.6% of the splits, it is in the top 3 most important features. Unfortunately, it is not very interpretable. The feature is calculated by fitting an autoregressive model to the timeseries, and evaluating its performance on 25 uniformly selected subsets of the timeseries of 10% the total length of the time series. The evaluation is based on 1-step ahead prediction. The difference between the real and predicted value forms a new TS, of length 192 in our case. The autocorrelation at lag 1 is calculated of each of these 25 TS. Finally, we take the absolute value of the mean of these 25 autocorrelation values. Further research could be done to determine why this particular feature is found to be this important. Other high-ranking features include those that quantify the level of surprise of a data point, given its recent memory. The remaining features show no clear pattern of type. As was the case for APB, we find that these lower ranked features’ prominence drops off rapidly.

The distributions for the most important features for each anatomy are shown in Figure 6. These figures were generated using kernel density estimation with a Gaussian kernel. Despite significant overlap in the distributions for the two classes, there is a definite difference between the two. For APB the distributions suggest that patients that are going to worsen have a more rapid return to an average of zero after the initial peak than patients that will not worsen. For AH such an intuitive interpretation is difficult due to the oblique nature of its most important TS feature. The features that were found have a few hyperparameters associated with them that could be optimized to further boost the performance of the classifier. We will not be doing this here, as these features were selected by looking at all splits at once, which covers the complete dataset. Their performance should be evaluated on another independent test set.

**Figure 6.**
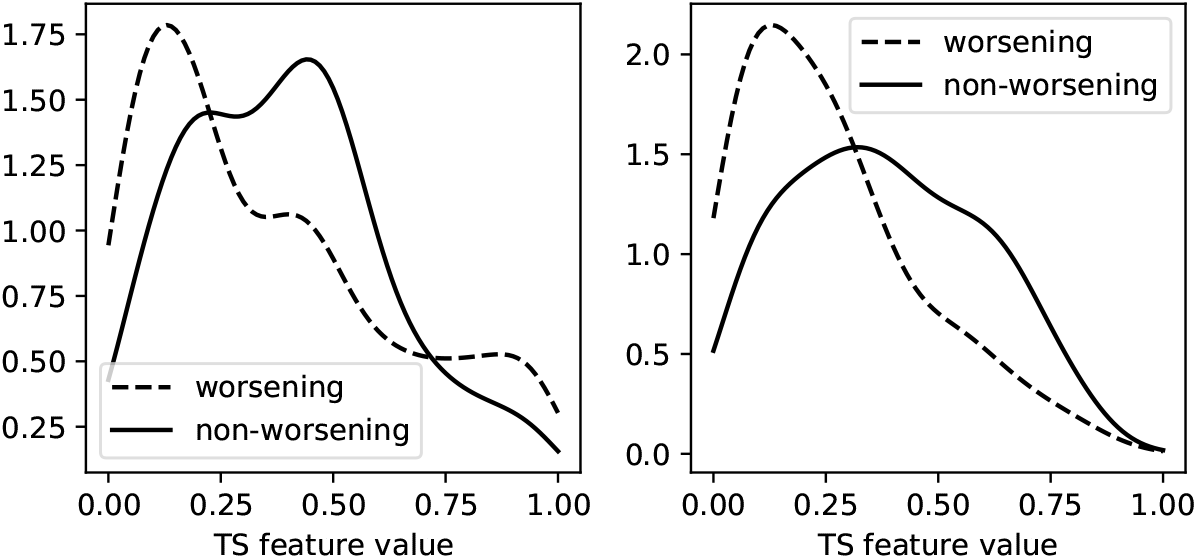
TS feature distributions. The distributions of the most important additional TS feature for APB (left) and AH (right). The dashed lines represent the distributions of the TS feature of patients that show progression after 2 years, whereas the solid lines are for those that do not progress. The distributions are normalized separately.

## 4 Discussion

This paper presents the first analysis on a new dataset, containing the full time series of several EP types. The idea was to extract a large number of features from the MEP from different time series analysis methods, and use a machine learning approach to see which ones are relevant. Improving the prediction of disability progression compared to using only the latencies, age, and EDSS at *T*_0_ was quite difficult, as the dataset seemed quite noisy, despite its larger size compared with the literature. On average, one in four TS features that remained after the mutual information and hierarchical clustering steps was found to contain at least some information relevant to the prediction task, though only a small subset contained a strong enough signal to be consistently marked important across multiple train/test splits. Nevertheless, a significant improvement was found by adding extra features that showed high importance. The usefulness of non-linear methods is also clearly demonstrated.

Much more remains to be investigated on this dataset. Given the large amount of literature on the usefulness of mmEP (as discussed in the introduction), the largest performance improvement is most likely achieved by including the VEP and SEP. Given the large differences in measurement times and frequencies of the different EP modalities, one has to decide between throwing away a lot of data, or using more elaborate techniques that robustly handle missing data. The second option that has potential for significant improvement is analyzing the whole longitudinal trajectory of the patient. This in contrast to our current analysis, where a single visit is used for predicting progression over 2 years. Inclusion of all (sparsely measured) mmEP and longitudinal modeling can be combined, and is an active research area [49, 50]. An obvious extension is to use TS algorithms not included in HCTSA. For example, another library with qualitatively different TS analysis methods is HIVE-COTE [51].

We have constricted ourselves to predicting progression over 2 years. This choice was made because it frequently occurs in the literature, and it leads to many training samples. Longer or shorter time differences are also of interest. It is, furthermore, believed by some clinicians that EPTS pick up disease progression faster than EDSS. One could check this by using short time-scale EPTS changes (e.g., 6 months) to predict EDSS changes on longer time-scales [22].

The obvious left-right-symmetry of the limb measurements is taken into account in a rudimentary way. Incorporating this symmetry in a more advanced way could boost performance. Data augmentation can be used to expand the size of the training set, which could stabilize the performance estimate. We note that even small neural networks are difficult to train on the current dataset. Data augmentation could make them competitive.

We performed a univariate TS analysis: only the EP with the maximal peak-to-peak amplitude is selected for feature extraction. Including all EPTS from each visit could provide extra information. Several algorithms that can handle multivariate TS are available, see e.g. [52]. However, the number of measured EPTS varies between visits and between limbs of the same visit. Dealing with this presents a technical challenge. We remark that our results point to the need for a larger number of samples if the dimensionality and redundancy of the input space is significantly increased, which would be the case for a multivariate TS analysis.

While the achieved AUC of 0.75 *±* 0.07 is impressive for a model with only MEP, EDSS at *T*_0_, age, and a few additional TS features, there is surely an upper limit to what mmEP can predict. Other variables such as, e.g., MRI, cerebrospinal fluid, and genomic data could boost performance [3]. A very important variable that is currently not included is the type of medication the patient is on. In the absence of a single, highly predictive marker, personalization will depend on a combinations of markers. Indeed, several studies show that a multi-parametric approach may improve our prognostic ability in MS [53, 54]. It involves the development of predictive models involving the integration of clinical and biological data with an understanding of the impact of disease on the lives of individual patients [55]. Besides the inclusion of extra biomarkers, another step of great practical importance is to move towards multicenter design studies. How well mmEP data from different centers can be combined remains an open and very important question [56, 57].

## 5 Conclusions

Multiple sclerosis is a chronic disease affecting millions of people worldwide. Gaining insight into its progression in patients is an important step in the process of gaining a better understanding of this condition. Evoked potential time series (EPTS) are one of the tools clinicians use to estimate progression. The prediction of disability progression from EPTS can be used to support a clinician’s decision making process regarding further treatment.

We presented a prediction model trained on a dataset containing all available EP measurements from the Rehabilitation & MS Center in Overpelt, Belgium. Any patient with two-year follow-up is included. It is an order of magnitude larger than most datasets used in previous works, and for the first time includes the raw time series, as opposed to just the high-level features extracted from them (i.e. latencies, peak-to-peak amplitude, morphology, etc.). The dataset consists of individuals undergoing treatment, which is clinically the most relevant scenario. We plan to make this dataset publicly available in the near future.

We found that adding additional features extracted from the raw time series improves performance, albeit marginally (∆AUC = 0.02 for the best performing classifier). Results suggest that the model would benefit from an increased dataset size. We found that linear models often used in previous works are significantly outperformed by the random forest classifier, especially when not adding extra TS features (∆AUC = 0.06). Given the limited number of biomarkers in the model (EDSS at *T*_0_, MEP, and age) and heterogeneity of the cohort, the reported performance (AUC 0.75 *±* 0.07) is quite good. We took an initial look at the features that were found to boost predictive power and found a few candidates that might be a good starting point for further research.

## Supporting information

Supplemental file 1: Top 20 features

## 6 List of abbreviations

A list of the abbreviations used throughout this work:

AH: Abductor Hallucis
AIC: Akaike Information Criterion
APB: Abductor Pollicis Brevis
AUC: Area Under Curve (of ROC, see below)
BAEP: Brainstem Auditory EP (EP see below)
BIC: Bayesian Information Criterion
EDSS: Expanded Disability Status Scale
EP: Evoked Potentials
EPS: EP Score
EPTS: EP Time Series
FWO: Fonds Wetenschappelijk Onderzoek
HCTSA: Highly Comparative Time-Series Analysis
LR: Logistic Regression
mmEP: Multimodal EP
MEP: Motor EP
MRI: Magnetic Resonance Imaging
MS: Multiple Sclerosis
MSE: Mean-Squared Error
PPMS: Primary Progressive MS
RF: Random Forest
ROC: Receiver Operating Characteristic
SD: Standard Deviation
SEP: Somatosensory EP
SPMS: Secondary Progressive MS
TS: Time Series
VEP: Visual EP

## Declarations

### Ethics approval and consent to participate

This study was approved by the ethical commission of the University of Hasselt (CME2017/729). No consent to participate/publish was necessary since this study uses retrospective data only.

### Consent for publication

Not applicable.

### Availability of data and materials

The dataset analyzed during the current study is currently not publicly available due to privacy concerns but is available from the corresponding author on reasonable request. We aim to release the dataset as well as the code used to generate the results publicly after taking steps to ensure the full anonymity of the patients in accordance with local data protection laws.

### Author’s contributions

JY performed the data analysis. JY, TB, DV, and LMP decided on the data analysis methodology. NH, VP and BVW provided clinical feedback for the data analysis. LMP coordinated the study. JY, TB, and LMP wrote the original manuscript. All authors read and approved the final manuscript.

## Acknowledgements

JY and TB are very grateful to the late Christian Van den Broeck for giving us the opportunity to work on this problem. The authors would like to thank Jori Liesenborgs and Geert Jan Bex for their valuable suggestions regarding the practical implementation of the data analysis pipeline, as well as Henny Strackx for his help during the data extraction phase at Overpelt.

## Additional Files

Additional file 1 — Most important TS features

This file contains tables of the 20 most prominent TS features across the 1000 train/test splits, for both APB and AH anatomies. It also provides a way of obtaining the code used to generate these features.

