## Supplemental file 1: Top 20 features for "Machine Learning Analysis of Motor Evoked Potential Time Series to Predict Disability Progression in Multiple Sclerosis"

March 2019

We provide the top 20 most important features for the disability progression task, both for APB (Table 1) and AH (Table 2). Their ranking is derived as follows: For each split we have 10 ranked features. For each unique feature that occurs across the splits we assign a score from 0 to 9 based on its position in a particular split (0 for being in first place, 9 for being in 10th). If it does not occur in a split it receives a score of 10. We add the scores of each of these features for all splits. The feature with the lowest score will be considered the most important. This score is included in the tables in the column *score*.

In the tables we also included the HCTSA ID, which can be used to get a description of the feature, as well as retrieve the code used to generate this feature. Instructions on how to do this can be found online[1].

The remaining columns are the name of the feature (as generated by HCTSA) and the percentage of splits where the feature occurs in the top $n$.

[1] `https://hctsa-users.gitbook.io/hctsa-manual/analyzing_visualizing/interpreting-features`

| name | hctsa id | top 1 | top 3 | top 5 | top 10 | score |
|---|---|---|---|---|---|---|
| SY SlidingWindow m s2 2 | 561 | 43.4% | 74.7% | 80.7% | 83.9% | 2414 |
| SY LocalGlobal l500.absmean | 762 | 3.8% | 26.0% | 50.5% | 82.8% | 5027 |
| CO AddNoise 1 gaussian.ami at 5 | 1264 | 2.4% | 24.1% | 50.5% | 79.4% | 5174 |
| SY SlidingWindow m s 4 1 | 554 | 4.4% | 26.7% | 42.8% | 59.6% | 6062 |
| SP Summaries pgram hamm.fpoly2 sse | 4307 | 14.5% | 38.2% | 43.4% | 45.3% | 6078 |
| PP Compare poly2.swms10 1 | 5795 | 3.6% | 17.2% | 32.5% | 51.4% | 6804 |
| PP Compare spline44.statav4 | 5912 | 15.9% | 27.0% | 30.7% | 33.3% | 7128 |
| PP Compare spline24.statav6 | 5882 | 0.3% | 5.8% | 15.8% | 43.6% | 8008 |
| SP Summaries pgram hamm.peakPower 2 | 4271 | 0.9% | 4.3% | 14.1% | 42.0% | 8163 |
| SY TISEAN nstat z 4 1 3.mean | 4778 | 4.8% | 12.1% | 16.0% | 19.5% | 8507 |
| SY SlidingWindow s ent2 10 | 597 | 0.1% | 0.8% | 5.3% | 31.0% | 8915 |
| SP Summaries pgram hamm.fpoly2csS p1 | 4304 | 0.3% | 2.6% | 8.6% | 19.6% | 9049 |
| PP Compare spline64.statav4 | 5943 | 0.8% | 5.2% | 8.5% | 14.1% | 9127 |
| CO Embed2 tau.eucds1 | 1928 | 0.1% | 0.9% | 4.0% | 20.5% | 9259 |
| SY TISEAN nstat z 4 1 3.std | 4783 | 0.2% | 2.3% | 5.8% | 15.6% | 9272 |
| FC Surprise T1 100 5 udq 500.uq | 2405 | 0.1% | 2.1% | 6.2% | 13.1% | 9316 |
| SP Summaries pgram hamm.fpoly2csS p2 | 4305 | 0.0% | 1.3% | 4.6% | 15.7% | 9325 |
| PP Compare medianf3.swms2 2 | 6040 | 0.1% | 1.9% | 5.5% | 14.1% | 9337 |
| MF AR arcov 4.res AC1 | 3861 | 0.0% | 0.5% | 3.1% | 19.0% | 9340 |
| SY SpreadRandomLocal 100 100.stdstd | 2136 | 0.0% | 1.0% | 4.4% | 16.5% | 9353 |

Table 1: The 20 most important features across all 1000 train/test splits for APB

| name | hctsa id | top 1 | top 3 | top 5 | top 10 | score |
|---|---|---|---|---|---|---|
| MF CompareTestSets y ar best uniform 25 01 1.ac1s mean | 7062 | 69.3% | 90.6% | 94.4% | 97.5% | 861 |
| SB BinaryStats iqr.longstretch0 | 3490 | 5.8% | 36.8% | 51.2% | 61.3% | 5420 |
| PH ForcePotential sine 1 1 1.median | 1655 | 1.6% | 14.9% | 31.9% | 53.7% | 6886 |
| FC Surprise T2 20 2 q 500.std | 2310 | 0.7% | 17.9% | 31.5% | 47.3% | 7106 |
| FC Surprise dist 20 2 q 500.mean | 2242 | 0.4% | 8.9% | 25.0% | 52.2% | 7297 |
| EN Randomize statdist.statav5diff | 2854 | 4.3% | 19.1% | 27.9% | 38.2% | 7357 |
| MF CompareTestSets y ar 4 rand 25 01 1.ac1s mean | 7042 | 1.5% | 16.5% | 28.5% | 39.9% | 7394 |
| WL DetailCoeffs db3 max.max1on2 median | 6526 | 0.9% | 10.5% | 21.1% | 31.3% | 8052 |
| FC Surprise dist 50 3 q 500.mean | 2250 | 0.0% | 1.8% | 8.8% | 31.1% | 8704 |
| FC Surprise dist 50 3 q 500.std | 2254 | 0.2% | 3.9% | 8.8% | 23.4% | 8927 |
| MF FitSubsegments arsbc uniform 25 01.sbcs range | 6981 | 0.0% | 3.8% | 7.8% | 22.8% | 8980 |
| EN Randomize statdist.statav5hp | 2855 | 2.5% | 8.0% | 10.6% | 13.7% | 8989 |
| EN Randomize permute.statav5fexpr2 | 2978 | 0.2% | 2.4% | 6.0% | 21.4% | 9050 |
| MF GP hyperparameters covSEiso covNoise 1 50 random i.meanS | 6396 | 1.1% | 5.0% | 8.7% | 12.4% | 9160 |
| MF GP hyperparameters covSEiso covNoise 1 50 random i.mlikelihood | 6389 | 0.8% | 6.0% | 8.4% | 10.6% | 9261 |
| FC LocalSimple mean3.tauresrat | 3017 | 3.7% | 6.6% | 7.0% | 7.4% | 9334 |
| SP Summaries pgram hamm.linfitsemilog all a2 | 4340 | 0.2% | 3.9% | 6.2% | 10.4% | 9376 |
| StatAvl500 | 550 | 0.2% | 1.6% | 4.9% | 12.5% | 9430 |
| SP Summaries fft logdev.linfitsemilog all a1 | 4587 | 0.0% | 1.7% | 4.6% | 12.1% | 9452 |
| FC LocalSimple lfittau.tauresrat | 3137 | 0.1% | 1.7% | 4.7% | 10.7% | 9481 |

Table 2: The 20 most important features across all 1000 train/test splits for AH
